# *Chlamydia trachomatis* effectors target the mitochondria and alter mitochondrial protein composition

**DOI:** 10.1101/2022.08.30.505961

**Authors:** Zoe Dimond, Laura D. Bauler, Yixiang Zhang, Aaron Carmody, Ted Hackstadt

**Author notes:** Preliminary accounts of this work were presented at the Gordon Conference on Microbial Toxins and Pathogenesis, 2010; and the Chlamydia Basic Research Society Biannual Meetings, 2011 and 2013.

## Abstract

Mitochondria are critical cellular organelles that perform a wide variety of functions including energy production and immune regulation. To perform these functions, mitochondria contain approximately 1,500 proteins, the majority of which are encoded in the nuclear genome, translated in the cytoplasm, and translocated to the mitochondria using distinct mitochondria targeting sequences (MTS). Bacterial proteins can also contain MTS and localize to the mitochondria. For the obligate intracellular human pathogen, *Chlamydia trachomatis*, interaction with various host cell organelles promotes intracellular replication. However, the extent and mechanisms through which *Chlamydia* interact directly with mitochondria remain unclear. We investigated the presence of MTS in the *C. trachomatis* genome and discovered 30 genes with around 70% or greater probability of mitochondrial localization. Five are translocated to the mitochondria upon ectopic expression in HeLa cells. Mass spectrometry of isolated mitochondria from infected cells revealed that two of these proteins localize to the mitochondria during infection. Comparison of mitochondria from infected and uninfected cells suggests that chlamydial infection affects mitochondrial protein composition. Around 125 host proteins were significantly decreased or absent in mitochondria from infected cells. Among these are pro-apoptotic factors and those related to mitochondrial fission/fusion dynamics. Conversely, 82 host proteins were increased in or specific to mitochondria of infected cells, many of which act as anti-apoptotic factors and upregulators of cellular metabolism. These data support the notion that *C. trachomatis* specifically targets host mitochondria to manipulate cell fate decisions and metabolic function to support pathogen survival and replication.

**Importance:** Obligate intracellular bacteria have evolved multiple means to promote their intracellular survival and replication within the otherwise harsh environment of the eukaryotic cell. Nutrient acquisition and avoidance of cellular defense mechanisms are critical to an intracellular lifestyle. Mitochondria are critical organelles that produce energy in the form of ATP and regulate programmed cell death responses to invasive pathogenic microbes. Cell death prior to completion of replication would be detrimental to the pathogen. *C. trachomatis* produces at least two and possibly more proteins that target the mitochondria. Collectively, *C. trachomatis* infection modulates mitochondrial protein composition favoring a profile suggestive of down-regulation of apoptosis.

## Introduction

The obligate intracellular pathogen, *Chlamydia trachomatis*, is the etiological agent of the most commonly reported bacterial infection, causing both blinding trachoma and sexually transmitted disease, affecting over 1.8 million people and 100 million people, respectively [1, 2]. *Chlamydia* have a unique biphasic developmental cycle where the bacteria alternate between two morphologically and functionally distinct forms; the infectious elementary body (EB) and the replicative reticulate body (RB). Critical to the developmental cycle is the establishment and maintenance of the chlamydia-containing vacuole, or inclusion. The inclusion serves as a protective membrane allowing for survival of the replicative body and as such, is a key interaction point between bacteria and host [3].

For *C. trachomatis*, inclusion interaction with host cells organelles has been demonstrated to be critical for nutrient uptake, inclusion localization and regulating escape mechanisms [4]. These known interactions occur through recruitment of organelles to the inclusion membrane through protein-protein interactions between inclusion membrane proteins (Incs) and organelle proteins. An example of these interactions includes the recruitment of Golgi-derived vesicles as a strategy for lipid acquisition from the host [5, 6], interactions of the inclusion membrane with the endoplasmic reticulum [7, 8], endocytic vesicles [9], lipid droplets [10], peroxisomes [11] and cytoskeleton [12]. The ability to manipulate mitochondria is critical for pathogens relying on the host for nutrients and survival, as is expected for the obligate intracellular pathogen *C. trachomatis*.

Mitochondria are critical cellular organelles with roles in diverse pathways including energy and metabolite production as well as regulation of cell survival. Often referred to as “the powerhouses of the cell,” these double membrane-bound structures are responsible for the production of most of cell’s ATP through the tricarboxylic acid (TCA) cycle and oxidative phosphorylation [13]. Along with production of ATP, these pathways are responsible for producing precursors for biosynthetic pathways, particularly fatty acid, nucleotide, and amino acid metabolism [14]. Inherently linked with mitochondria’s role in energy production is the maintenance of homeostasis within the cell, and therefore the promotion of cell death in dysregulated states, such as during stress or infection. Intrinsic apoptosis is induced when an internal stimulus leads to the activation of pro-apoptotic factors, which results in the depolarization and permeabilization of the outer mitochondrial membrane, ultimately leading to cell death [15, 16].

Due to their diverse functions, mitochondria contain many proteins functioning in structural and enzymatic roles. Mitochondrial DNA encodes only about 1% of the mitochondrial proteome, most of which are important in oxidative phosphorylation or translational machinery [17]. However, hundreds of proteins found within the mitochondria are translated in the cytoplasm and translocated to the mitochondria using distinct <u>m</u>itochondria <u>t</u>argeting <u>s</u>equences (MTS). These MTS are typically amphiphilic cleavable signals between 20 and 40 amino acids in length on the N-terminus containing multiple positively charged residues and no acidic residues [18]. It has been shown that bacterial proteins can also contain MTS and localize to the mitochondria [19, 20]. For example, enteropathogenic *Escherichia coli* protein EspF is injected into the host via a Type 3 secretion system (T3SS) and localizes to the mitochondria, causing cytochrome c release and apoptosis [21].

While it has been shown that *Chlamydia* modulate host processes associated with mitochondria, such as inhibition of apoptosis and promotion of mitochondrial fusion [22], the extent and mechanisms through which *Chlamydia* interact directly with mitochondria remain unclear. Interestingly, mitochondria are not recruited to the *C. trachomatis* inclusion membrane, like other organelles [23]. We hypothesized that rather than recruiting mitochondria to the inclusion, secreted effectors might instead be trafficked to the mitochondria. In this study, we show that *C. trachomatis* encodes proteins which contain MTS, are Type 3 secreted and localize to the mitochondria during infection. Additionally, mitochondria from infected cells have unique proteomes which reflect global changes caused by the presence of *C. trachomatis*.

## Methods

### Strains and Cell Culture

*C. trachomatis* serovar L2 (LGV 434/Bu) was propagated in HeLa 229 cells in RPMI 1640 media containing 5% FBS at 37 °C with 5% CO_2_. EBs were purified by Renografin density gradient as previously described [24] and infectious EBs were quantified by titration and manual counting as described [25] using indirect immunofluorescence with a rabbit polyclonal anti-*C. trachomatis* L2 EB antibody followed by an anti-rabbit IgG AlexaFluor 488-conjugated secondary antibody (Invitrogen). Chlamydial genomic DNA was purified using the DNeasy Blood and Tissue Kit (Qiagen). Briefly, purified EBs were boiled for 10 minutes then incubated at room temperature in DTT at a final concentration of 20% v/v for 15 minutes. The DNeasy Blood and Tissue DNA isolation kit (Qiagen) was used to extract genomic DNA following manufacturer’s instructions for Gram-negative bacteria.

### Generation of CT-GFP fusion constructs

For each candidate effector, the open reading frame was amplified from *C. trachomatis* L2 genomic DNA by PCR with primers that added restriction sites on each end using AccuPrime Pfx DNA Polymerase (Invitrogen). DNA was extracted by gel purification using the Qiaquick Gel Extraction Kit (Qiagen). The pEGFP-N1 plasmid was purified using the MaxiPrep kit (Qiagen). Plasmid and insert products were digested using appropriate restriction enzymes (New England Biolabs) and ligated using T4 Ligase (New England Biolabs). Constructs were transformed into chemically competent Turbo *E. coli* (New England Biolabs) as per manufacturer instructions and transformants verified by colony PCR and Sanger sequencing. Plasmids were extracted using a MaxiPrep kit. For specific effectors, the MTS was removed by generating additional primers inside the predicted MTS (or the first 33 amino acids for CT642) and adding the start codon back into the forward primer sequences.

### Transfections and Microscopy

Purified plasmids were transfected into HeLa 229 cells using the Xfect transfection reagent (Takara Biosciences) according to manufacturer’s instructions. Briefly, 10^5^ cells were plated onto 24-well plates with glass coverslips and allowed to adhere overnight. Plasmid DNA (1ug per well) was mixed with reaction buffer (up to 25 uL per well). Polymer (0.3 uL per well) was mixed with reaction buffer (up to 25uL per well), added to the plasmid-reaction buffer solution and incubated at room temperature for 10 minutes. 50 uL of solution was added to each well and incubated for 4 hours before the media was replaced with fresh RPMI plus 5% FBS. Transfections were incubated 48 hours before staining with 100nM MitoTracker Red (Invitrogen) and DAPI (NucBlue Live ReadyProbes, Invitrogen) for 20 minutes at 37°C. Coverslips were washed twice in PBS and imaged under 60X magnification on a Nikon Ti2e microscope using DAPI, FITC (GFP) and TRITC (MitoTracker) filters. Colocalization was determined quantitatively by NIS-Elements 64-bit software (version 5.11.02, Nikon).

### Type 3 Secretion Analysis by *Yersinia pseudotuberculosis*

The first 50 or 100 amino acids of *C. trachomatis* genes were fused to Npt (Neomycin phosphotransferase) in the pFlag-CTC as previously described [26, 27]. The constructs were transformed into wildtype *Y. pseudotuberculosis* (pIB102) or the *ΔyscS* strain (pIB68) and cultures were grown at 26°C for 2 hours either in the presence or absence of 5mM calcium. Cultures were shifted to 37°C and protein fusion expression was induced with 0.01mM IPTG. Cultures were harvested after 4 hours post induction and separated into pellet or supernatant fractions. Samples were analyzed by immunoblot with anti-Flag M2 antibodies or YopN antibodies.

### Isolation of Mitochondria

In triplicate experiments, four T-75 flasks were seeded with 1x 10^6^ HeLa 229 cells and allowed to adhere overnight. Two flasks were infected with *C. trachomatis* L2 at a multiplicity of infection of 1 by centrifugation at 550 x g for 20 minutes at room temperature, fed with RPMI plus 5% FBS and incubated at 37°C for 20 hours. The remaining flasks were mock infected. After incubation, cells were trypsinized and resuspended in PBS. One infected and one uninfected flask were treated with DSS Crosslinker (Thermo Scientific) following manufacturer instructions for intracellular crosslinking. All samples were then resuspended in FACS buffer (PBS with 1% BSA and 2.5mM EDTA) [28] and stained with 100nM MitoView 405 (Invitrogen) for 15 minutes at 37°C. Cells were pelleted and resuspended in ice cold cell lysis buffer (200mM sucrose, 10mM Tris pH 7.4, 0.5mM EDTA and 1X Halt Protease Inhibitor Cocktail (Invitrogen) in PBS).

Mitochondria were sorted on a MACSQuant Tyto Cell Sorter (Miltenyi Biotech). The MACSQuant Tyto HS Cartridge (Miltenyi Biotec) was primed using 0.4 ml of MACSQuant Tyto Running Buffer (Miltenyi Biotec) according to manufacturer instructions. Fluorescently labeled beads were used to accurately identify the gating threshold for removal of debris and instrument noise. Cell lysates were loaded to a MACSQuant Tyto HS Cartridge (#130-121-549, Miltenyi Biotec) and sorted according to the instrument instructions until 1 x 10^7^ positive events had been collected. Purified mitochondria were resuspended with 2X Laemmli Buffer with β-mercaptoethanol and incubated at 100°C for 10 minutes to denature proteins.

### Sample preparation and LC-MS/MS

Mitochondrial proteins were denatured by 10 mM tris (2-carboxyethyl) phosphine (TCEP) for 45 min at 56 °C and alkylated by 20 mM iodoacetamide for 1 hr at room temperature in the dark. The SP3 protocol was applied for protein cleanup [29]. On-bead trypsin digestion was performed at 37 °C overnight. The resulting peptides were desalted with C18 Zip-tip (Millipore, Bedford, MA, USA) for LC-MS/MS analysis.

A Q Exactive plus mass spectrometer (Thermo Fisher Scientific, San Jose, CA, USA) coupled with an Easy-nLC 1200 HPLC system was utilized and controlled by Xcalibur software (Thermo Fisher Scientific). Peptide samples were loaded onto an Acclaim PepMap™ 100 C18 trap column (75 μm × 20 mm, 3 μm, 100 Å) in 0.1% formic acid and further separated on an Acclaim PepMapTM RSLC C18 analytical column (75 μm × 250 mm, 2 μm, 100 Å) using an acetonitrile-based gradient (Solvent A: 0% acetonitrile, 0.1% formic acid; Solvent B: 80% acetonitrile, 0.1% formic acid) at a flow rate of 300 nL/min. The gradient was as follows: 0-90 min, 2-25% B; 90-120 min, 25-40% B; 120-125 min, 40-100% B; 125-127 min, 100% B, followed by column wash and re-equilibration to 2% solvent B. Electrospray ionization was carried out with an EASY-Spray™ Source at a 275°C capillary temperature, 50 °C column temperature, and 1.9 kV spray voltage. The mass spectrometer was operated in data-dependent acquisition mode with mass range 350 to 2000 m/z. Full scan resolution was set to 70,000 with AGC target at 1e6 and a maximum fill time of 30 ms. Fragment scan resolution was set to 17,500 with AGC target at 5e4 and a maximum fill time of 50 ms. Normalized collision energy was set to 27. The dynamic exclusion was set with 60 s duration and a repeat count of 1.

### MS data analysis

Raw data were acquired by the Xcalibur 4.2 system (Thermo Scientific, Bremen, Germany) and analyzed by Proteome Discoverer 2.4. The raw files were searched against *Homo sapiens* (SwissProt v2017-10-25), *Chlamydia trachomatis* (UniProtKB UP000000795) and common contaminants provided by MaxQuant. Search parameters were as follows: full trypsin digestion with two missed cleavages. Carbamidomethylation on cysteine residues was set as the fixed modification. A total of five variable modifications were allowed per peptide from the following list: oxidation on methionine, acetylation on protein N-terminus, pyro-glutamate conversion on N-term glutamate, methionine-loss on protein N-terminus, and methionine-loss and acetylation on protein N-terminus methionine. The precursor peptide mass tolerance was set to 5 ppm. The fragment ion mass tolerance was set to 0.02 Da. The Percolator algorithm was used for PSM validation. Proteins reported in the result have at least two distinct peptides detected in the analysis. Proteins detected fewer than 2 times in 3 replicates were excluded from statistical analysis. The statistical analysis of these data was completed using unpaired, two-tailed student’s t test.

### Bioinformatic analyses

To predict MTS, open reading frames were translated from *C. trachomatis* serovar D (D/UW-3) and analyzed with MitoProt II V1.101 [30]. Proteins with export to mitochondria probability scores over 0.7 were selected as candidate genes. *C. trachomatis* L2 proteins from the proteomics dataset were also analyzed for MTS. Type 3 secretion prediction was performed using T3Sepp (accessible from http://www.szu-bioinf.org/T3SEpp/) [31]. Functional analysis of proteomics hits was performed using DAVID (<u>D</u>atabase for <u>A</u>nnotation, <u>V</u>isualization and <u>I</u>ntegrated <u>D</u>iscovery) Bioinformatics resources (accessible from https://david.ncifcrf.gov/home.jsp) [32, 33]. Graphs were generated using Graphpad Prism (version 9.3.1).

## Results

### *C. trachomatis* proteins contain predicted and functional mitochondria targeting sequences

To investigate the presence of MTS within the chlamydial genome, a bioinformatic screen was used to analyze the primary sequence for *C. trachomatis* D/UW-3 proteins. The computational MTS predictor, MitoProt [30], calculates a mitochondrial export probability based on N-terminal MTS parameters. Thirty *C. trachomatis* proteins with predicted MTS and mitochondrial export probabilities over or near 0.7 were identified (Table 1). The majority of candidate proteins were chlamydia-specific hypothetical proteins but five putative or demonstrated inclusion membrane proteins, two known Type 3 secreted proteins and two of the putative cytotoxin pseudogenes were represented as well.

**Table 1.** *C. trachomatis* proteins with mitochondria targeting sequences by MitoProt prediction

| Gene | Annotated function | MTS | Mitochondrial export probability | Mitochondrial localization? |
| --- | --- | --- | --- | --- |
| CT011 | Hypothetical Protein | 1:47 | 0.979 | No |
| CT037 | Hypothetical Protein | 1:70 | 0.742 | NT |
| CT058 | Hypothetical Protein | 1:87 | 0.985 | No |
| CT060 | FHIPEP family type III secretion protein | 1:154 | 0.969 | NT |
| CT080 | Late transcription unit protein B (ItuB) | 1:24 | 0.976 | No |
| CT087 | 4-alpha glucanotransferase (malQ) | 1:41 | 0.941 | No |
| CT101 | Hypothetical Protein | 1:70 | 0.858 | NT |
| CT132 | Hypothetical Protein | 1:34 | 0.962 | Yes |
| CT133 | Class I SAM-dependent methyltransferase | 1:17 | 0.831 | No |
| CT166 | Putative cytotoxin pseudogene | 1:35 | 0.945 | NT<sup>a</sup> |
| CT168 | Putative cytotoxin pseudogene | 1:29 | 0.810 | NT |
| CT195 | Putative inclusion membrane protein | 1:127 | 0.960 | No |
| CT229 | Inclusion membrane protein, CpoS | 1:97 | 0.712 | No |
| CT244 | Hypothetical Protein | 1:8 | 0.686 | No |
| CT256 | Hypothetical Protein | 1:155 | 0.757 | No |
| CT300 | Hypothetical Protein | 1:112 | 0.952 | No |
| CT339 | DNA internalization-related competence protein, ComEC | 1:135 | 0.997 | No |
| CT351 | Hypothetical Protein | 1:45 | 0.676 | NT |
| CT352 | Hypothetical Protein | 1:48 | 0.962 | NT |
| CT385 | Hypothetical Protein | 1:5 | 0.7054 | No |
| CT484 | Hypothetical Protein | 1:85 | 0.846 | No |
| CT529 | Putative inclusion membrane protein | 1:37 | 0.985 | Yes |
| CT550 | Hypothetical Protein | 1:3 | 0.887 | No |
| CT552 | Hypothetical Protein | 1:47 | 0.996 | No |
| CT616 | Hypothetical Protein | 1:86 | 0.968 | No |
| CT618 | Putative inclusion membrane protein | 1:37 | 0.896 | Yes |
| CT642 | Putative inclusion membrane protein | 1:8 | 0.808 | Yes |
| CT647 | Hypothetical Protein | 1:32 | 0.753 | Yes |
| CT700 | Hypothetical Protein | 1:40 | 0.717 | No |
| CT862 | Type III secretion low calcium response chaperone, IcrH2 | 1:47 | 0.735 | No |

<sup>a</sup> NT denotes Not Tested

Eukaryotic expression vectors were generated using the pEGFP-N1 plasmid to express each candidate gene as an EGFP protein fusion that could be expressed ectopically in HeLa cells. We utilized *C. trachomatis* L2 coding regions for the ability to perform future genetic experiments, and therefore were unable to generate vectors for CT166, CT168, and CT352 which are absent in the *C. trachomatis* L2 genome [34]. Genes with known functions (CT060 [35], CT101 [36] or pseudogenes CT037 and CT300) were excluded from construct generation while CT616 proved challenging to clone. The remaining twenty-two candidate genes from the MTS screen were examined for protein localization in HeLa cells (Table 1).

Five candidates, CT132, CT529, CT618, CT642 and CT647 showed colocalization with mitochondria upon transfection when compared with the diffuse cytoplasmic staining of the vector control (Figure 1A). Colocalization was quantified with Pearson correlation coefficients. Closer examination of the MTS for each of these proteins revealed that each was enriched for canonical amino acids; alanine, arginine, leucine, and serine as well as for positive residues (Figure 1B). CT529, CT618 and CT642 are all predicted inclusion membrane proteins with characteristic bilobed transmembrane domains, although CT529 has a possible third transmembrane domain at the C-terminus of the protein [37]. CT132 is a hypothetical protein with six putative transmembrane domains, which may be characteristic of a multipass integral membrane protein [38]. CT132 also shows homology, using the Basic Local Alignment Search Tool (BLAST), to transporter protein BrkB of *Bordetella pertussis* which similarly has six membrane-spanning domains [39]. CT647 is a hypothetical protein with no known or predicted functional domains.

**Figure 1.**
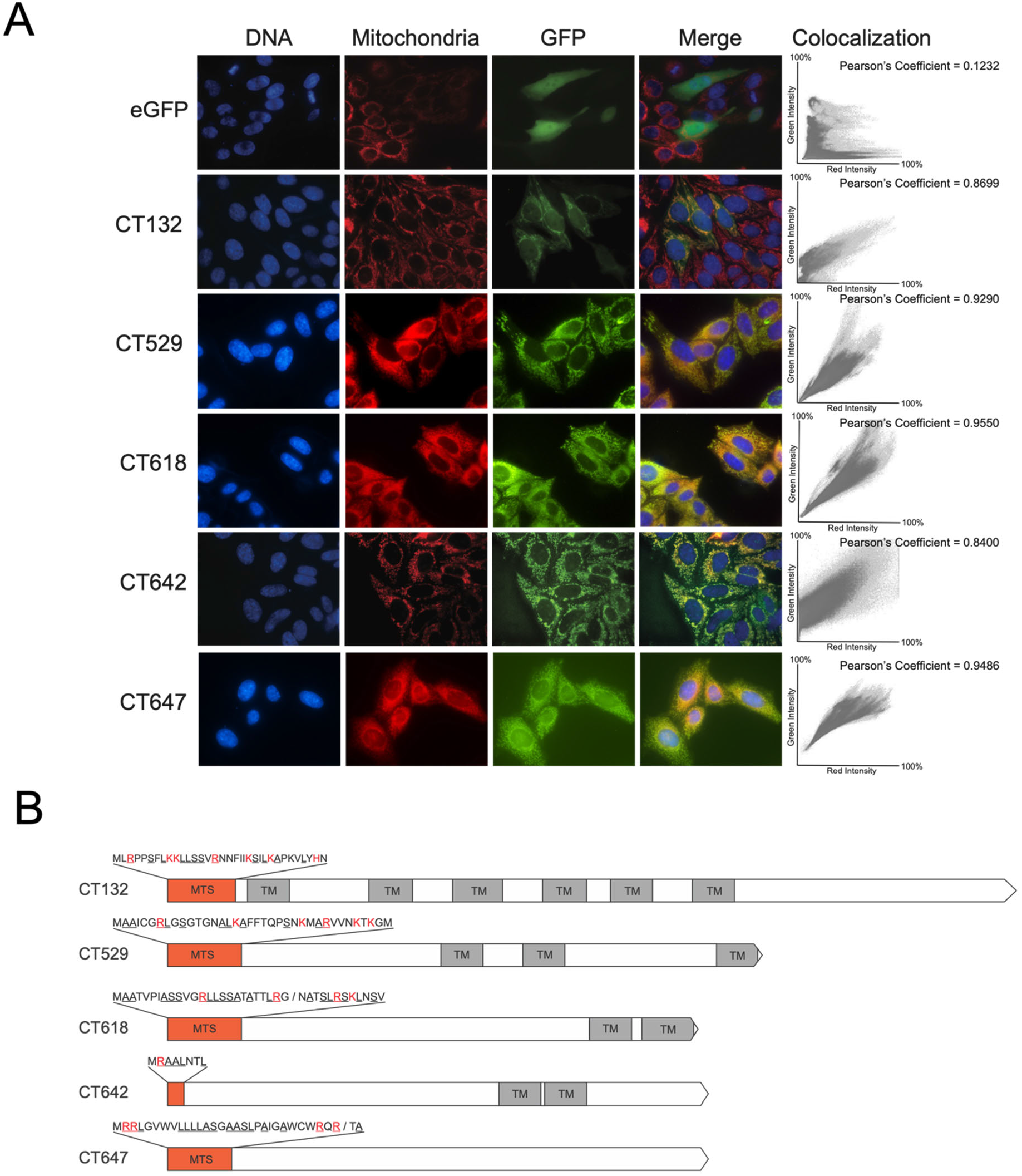
A) Immunofluorescent staining shows GFP localization of transfected *C. trachomatis* proteins with functional MTS. HeLa cells on coverslips were transfected with either the empty EGFP-N1 vector (eGFP) or GFP-tagged *C. trachomatis* proteins (CT###) and incubated for 48 hours before staining with 100nM MitoTracker (red) and nucBlue (blue). Bar = 10 μm. Coverslips were imaged under 60X magnification. Colocalization scatterplots were determined for the regions of interest in each representative image and Pearson coefficients are reported. B) Gene maps for the five candidate MTS sequences. Orange boxes represent MTS sequences, which are inset above. Red amino acids indicate positive residues and underlined residues denote canonically enriched amino acids. Predicted cleavage sites are indicated with a slash.

To verify that the mitochondrial localization for these five proteins was due to recognition of the MTS, constructs were designed where the N-terminal predicted MTS was removed, leaving the methionine start codon intact. For all five mitochondria-localized constructs, removal of the MTS resulted in an inability to colocalize with mitochondria (Figure 2). Both CT132-GFP and CT618-GFP signals were cytoplasmic and appeared similar to the majority of non-functional MTS proteins (Supplemental Figure 1). CT529, CT618 and CT647 GFP signals had punctate staining that was no longer correlated with mitochondria. These data suggest that the bioinformatically predicted MTS are functionally recognized in HeLa cells for CT132, CT529, CT618 and CT647, resulting in translocation of these proteins to the mitochondria.

**Figure 2.**
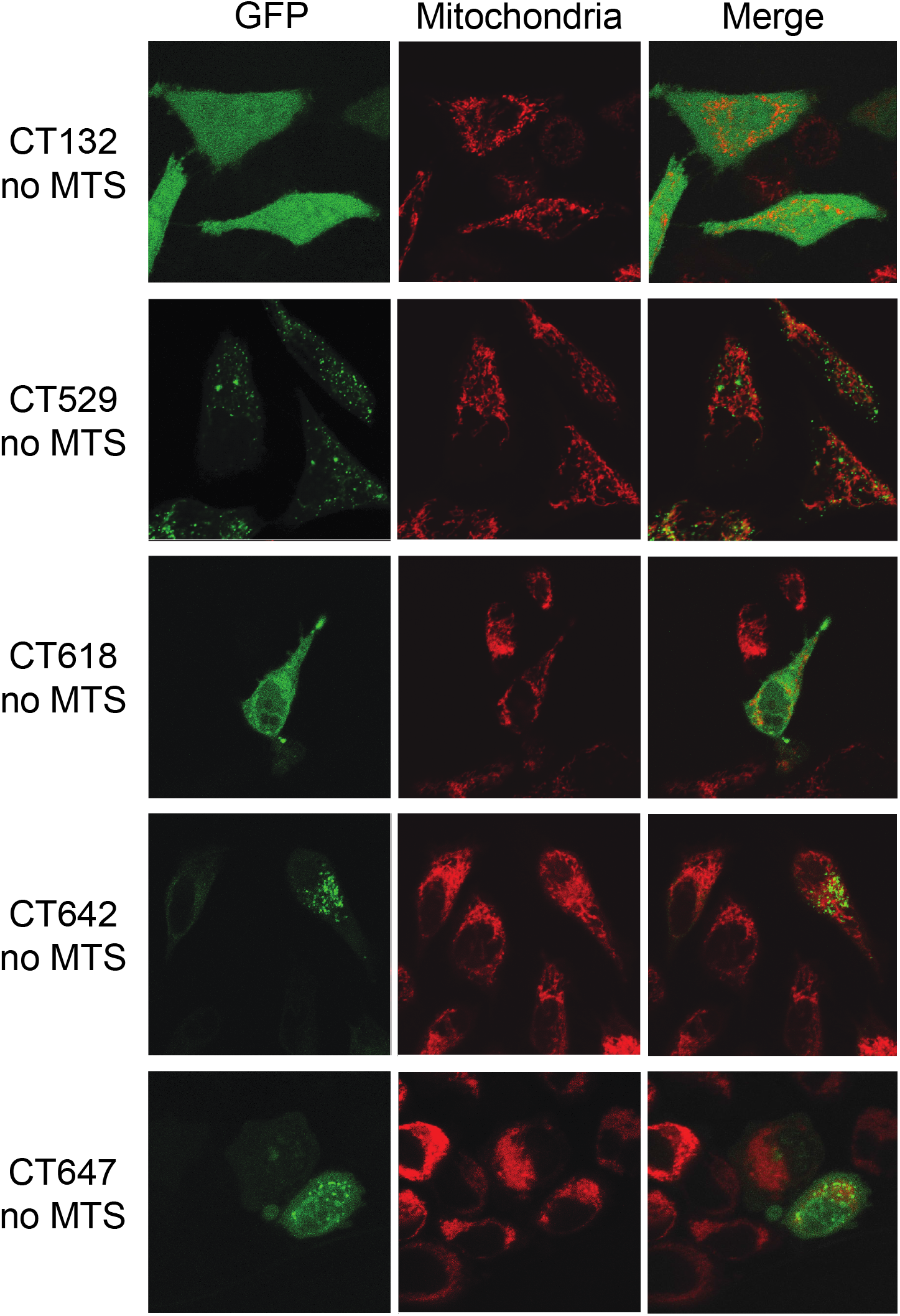
Immunofluorescent staining shows GFP localization of mitochondria-targeted *C. trachomatis* proteins after removal of the MTS. HeLa cells on coverslips were transfected with MTS removed *C. trachomatis* protein GFP fusions (CT### no MTS) and incubated for 24 hours before staining with 100nM MitoTracker (red). Bar = 10 μm.

Seventeen proteins did not localize with mitochondria after transfection (Supplemental Figure 1A). The majority of these candidates had diffuse cytoplasmic localization, similar to the EGFP vector control. Three protein fusions appeared to localize to discrete sites within the cell other than the mitochondria. CT229 is a known inclusion membrane protein, CpoS, and inserts into the inclusion membrane where it interacts with Rab GTPases to regulate vesicular trafficking [40]. The CT229-GFP fusion protein in this study appears to be maintaining interaction with clathrin-coated vesicles of the trans-Golgi network, as previously described [41]. Hypothetical protein CT700-GFP fusion staining was similar to CT229, and we demonstrate that this protein also localized to the Golgi apparatus during infection (Figure 1B). Additionally, putative inclusion membrane protein CT195 has previously been demonstrated to colocalize with endoplasmic reticulum (ER) structures and our construct had similar colocalization with calnexin stained ER membranes (Supplemental Figure 1B) [42].

### CT529, CT618 and CT642 are Type 3 secreted proteins

In order for the MTS to be recognized by cognate chaperones and transporters, the chlamydial MTS would need to be present in host cytosol, therefore, we examined if the five mitochondrial-targeting candidate proteins could be secreted. In *C. trachomatis*, the Type III secretion system (T3SS) is critical for the delivery of effectors into the host cell, as well as Incs into the inclusion membrane [43]. The bioinformatic prediction pipeline T3SEpp was used to predict likelihood of T3S for each protein coding sequence [31]. Both CT132 and CT647 were identified as non-T3S while CT529, CT618 and CT647 were predicted to be T3S, which matches their identification as putative Incs (Figure 3A).

**Figure 3.**
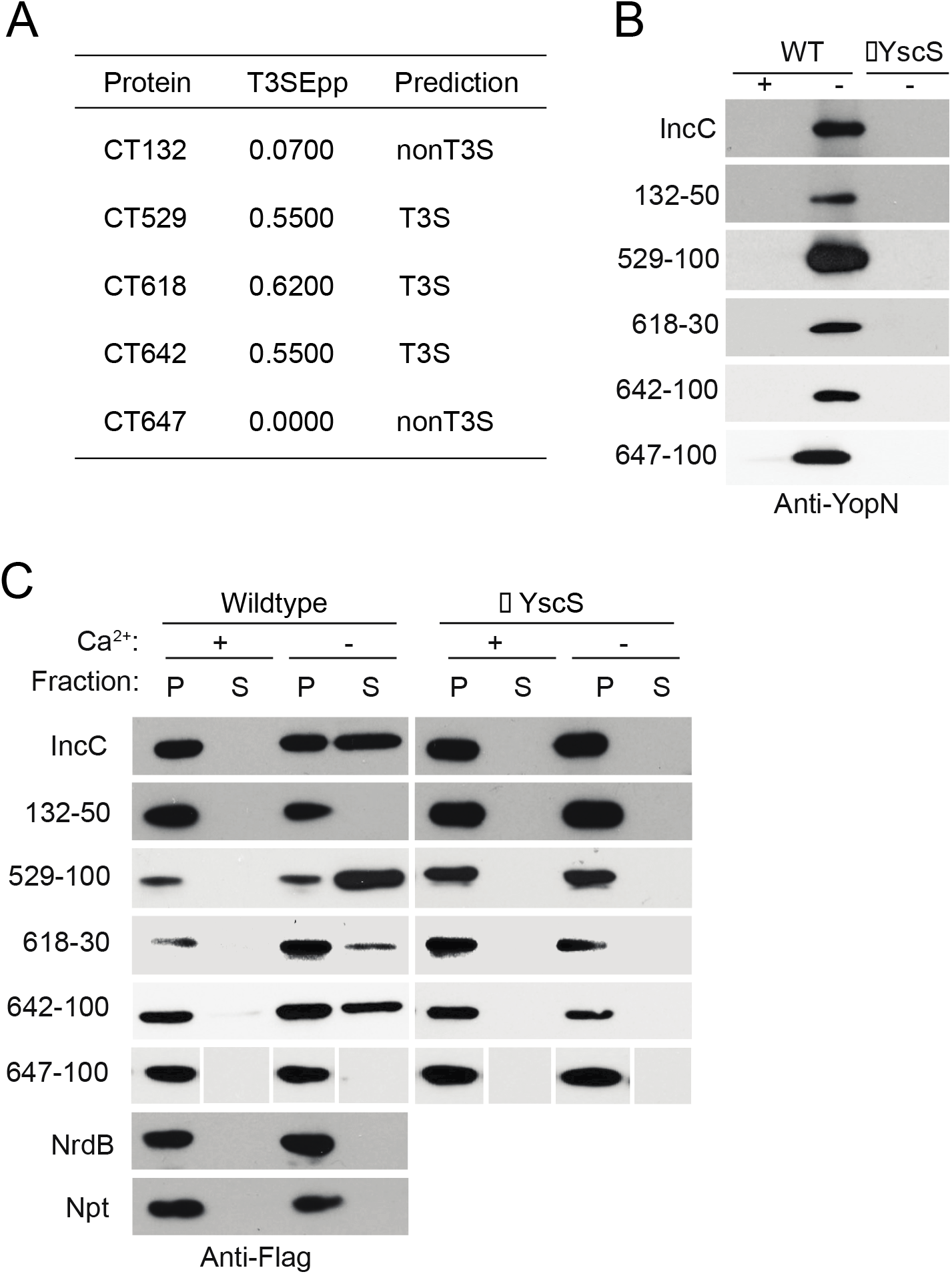
Select mitochondrial targeting *Chlamydia* proteins are secreted by Type III secretion. A) Results from computational integrated prediction pipeline, T3SEPP, predicting bacterial Type 3 effectors. B) *Y. pseudotuberculosis* expressing the N-terminal sequence (30-100bp) of chlamydial proteins, or the vector control were grown in the presence or absence of 5mM Ca2+, where the absence promotes Type 3 secretion. Cultures were induced with 0.01mM IPTG and temperature shifted from 26 to 37 degrees Celsius and supernatants were probed for YopN (*Yersinia* T3SS control). C) Flag-tagged Npt-fusion constructs were expressed as in B) and samples were fractionated into pelleted cells (P) for non-secreted fractions and supernatant (S) for secreted fractions. The samples from 647-100 were run in a different order and certain panels were spliced for consistency in the presentation of this figure as indicated by gaps.

The chlamydial T3SS is heterologous to that of *Yersinia pseudotuberculosis*, which can be utilized as a model to demonstrate T3S of *C. trachomatis* proteins [26, 27]. The N-terminal sequence of the five candidate effectors were fused to the T3S carrier protein neomycin phosphotransferase (Npt) of either wildtype *Y. pseudotuberculosis* or the T3SS-deficient ΔYscS strain. As a control, we probed for *Yersinia* secreted effector YopN in the supernatants for each of the samples to ensure that expression of the fusion construct did not disrupt normal yersinial T3S (Figure 3B). In wildtype *Y. pseudotuberculosis*, fusion of CT132 and CT647 N-terminal regions to Npt did not result in secretion in the absence of calcium, which activates T3S and were not different from the NrdB negative control which is a chlamydial gene with no T3S signal. In contrast, CT529, CT618 and CT642 fusion constructs were comparable to the IncC positive control which is a known T3S effector in *C. trachomatis* and has been previously demonstrated to be T3S in *Y. pseudotuberculosis* (Figure 3C) [27]. In the ΔYscS strain, these constructs were not detected in the supernatant, demonstrating that these are in fact Type 3 secreted effectors.

### CT529 and CT618 proteins are identified in mitochondria of infected cells

To confirm the localization of the candidate effectors during infection, a proteomics approach was taken. Mitochondria from HeLa cells either uninfected or infected for 24 hours with *C. trachomatis* L2 were isolated using fluorescence-activated mitochondrial sorting [28]. Isolated mitochondria were lysed and total protein content was identified using mass spectrometry. Triplicate experiments were performed for each condition with and without DSS crosslinking.

Mitochondrial proteins were either cross-linked or not, however, there was no significant difference between crosslinked and non-crosslinked conditions. No *C. trachomatis* proteins were present only in cross-linked conditions. Forty-nine chlamydial proteins with over two unique peptides in at least two replicates were detected in the mitochondria of infected cells after elimination of those detected in the uninfected condition. Of the original candidate proteins, both CT529 and CT618 were detected in at least four replicates (Table 2). CT132, CT642 and CT647 were not identified in any replicate. Of the remaining chlamydial hits, 12 (24.5%) of the hits were ribosomal proteins and five (10.2%) were hypothetical proteins.

**Table 2.**
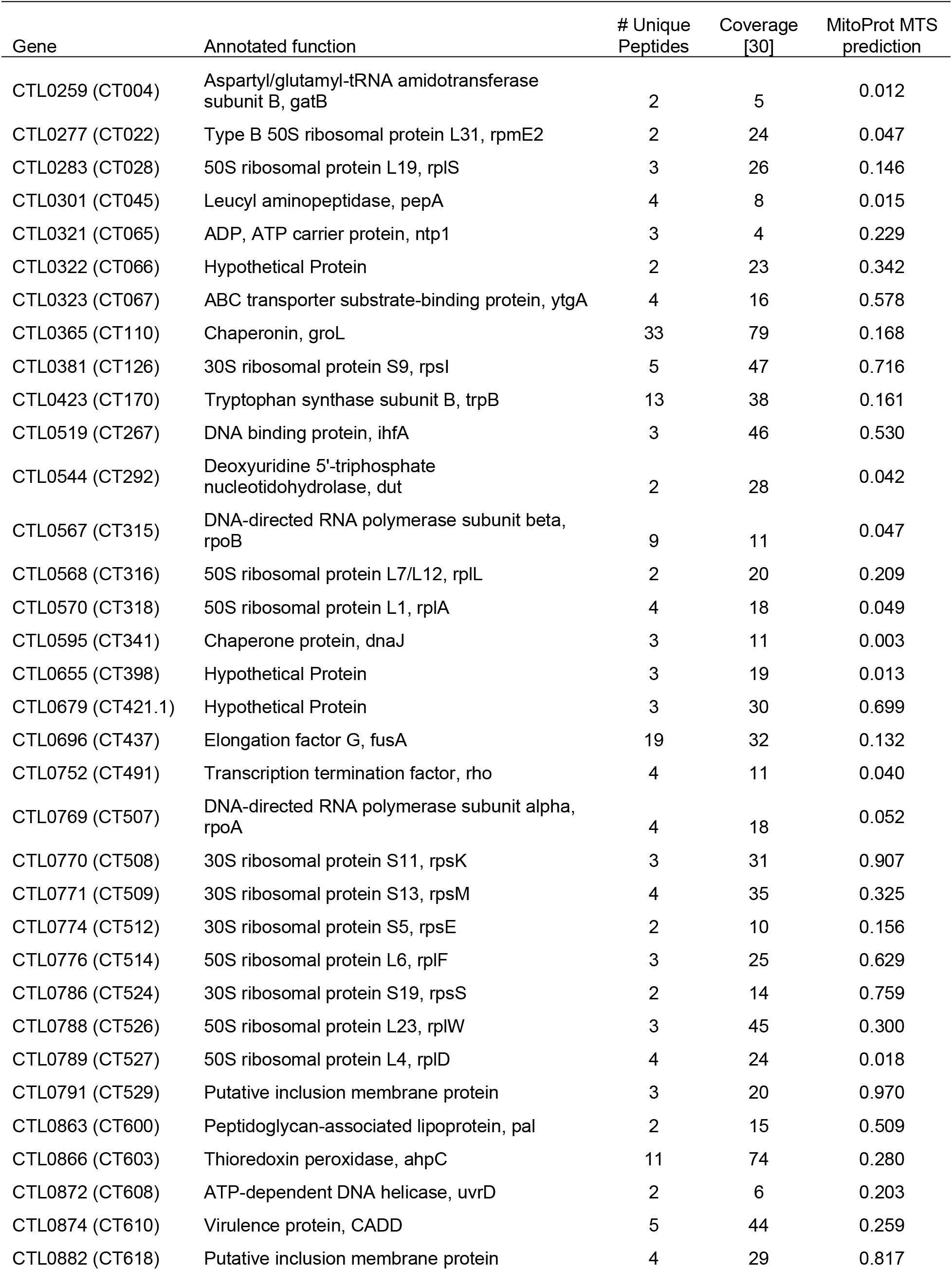
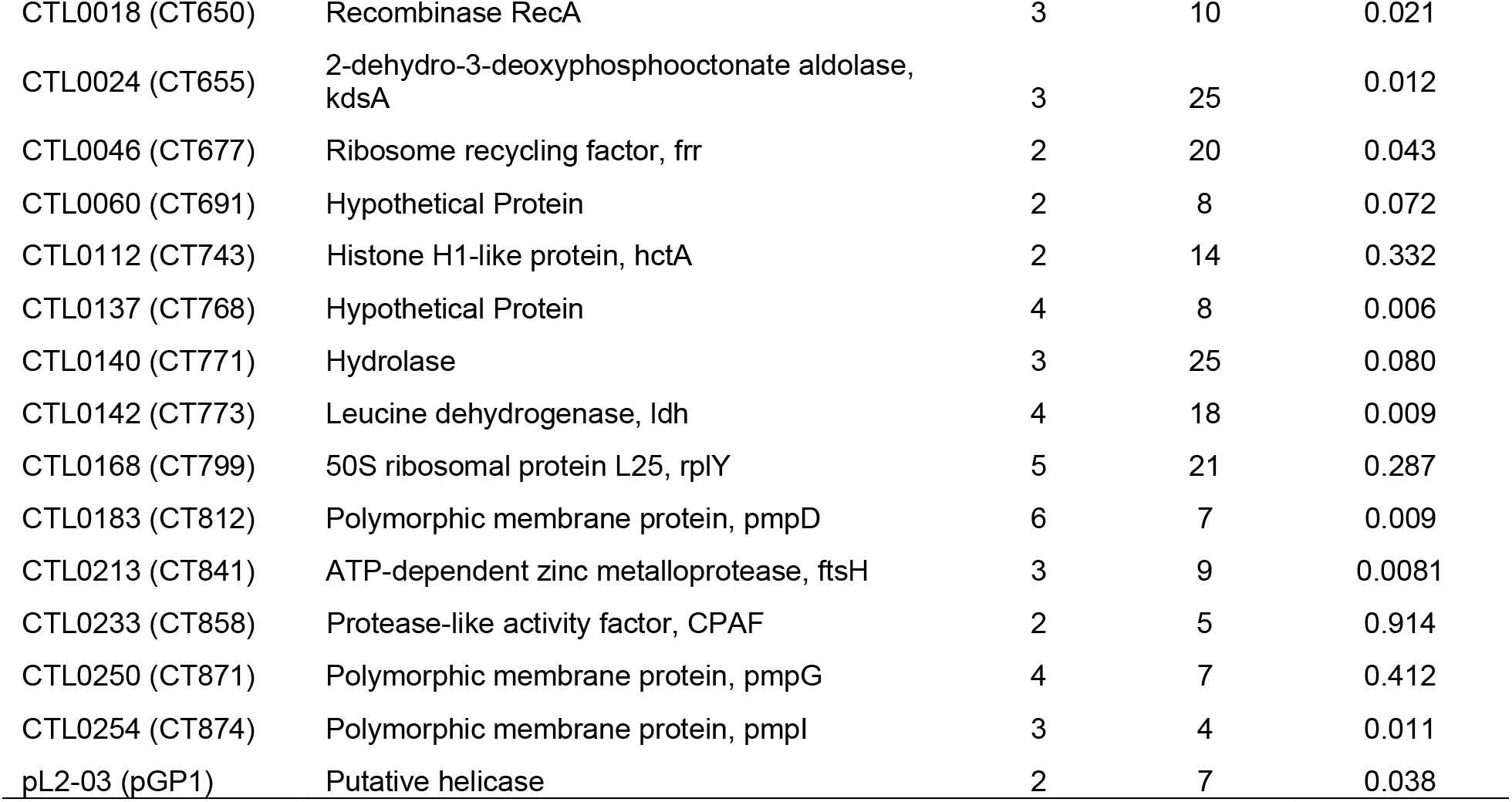
*C. trachomatis* proteins identified in mitochondria from infected cells

| Gene | Annotated function | # Unique Peptides | Coverage [30] | MitoProt MTS prediction |
| --- | --- | --- | --- | --- |
| CTL0259 (CT004) | Aspartyl/glutamyl-tRNA amidotransferase subunit B, gatB | 2 | 5 | 0.012 |
| CTL0277 (CT022) | Type B 50S ribosomal protein L31, rpmE2 | 2 | 24 | 0.047 |
| CTL0283 (CT028) | 50S ribosomal protein L19, rplS | 3 | 26 | 0.146 |
| CTL0301 (CT045) | Leucyl aminopeptidase, pepA | 4 | 8 | 0.015 |
| CTL0321 (CT065) | ADP, ATP carrier protein, ntp1 | 3 | 4 | 0.229 |
| CTL0322 (CT066) | Hypothetical Protein | 2 | 23 | 0.342 |
| CTL0323 (CT067) | ABC transporter substrate-binding protein, ytgA | 4 | 16 | 0.578 |
| CTL0365 (CT110) | Chaperonin, groL | 33 | 79 | 0.168 |
| CTL0381 (CT126) | 30S ribosomal protein S9, rpsI | 5 | 47 | 0.716 |
| CTL0423 (CT170) | Tryptophan synthase subunit B, trpB | 13 | 38 | 0.161 |
| CTL0519 (CT267) | DNA binding protein, ihfA | 3 | 46 | 0.530 |
| CTL0544 (CT292) | Deoxyuridine 5'-triphosphate nucleotidohydrolase, dut | 2 | 28 | 0.042 |
| CTL0567 (CT315) | DNA-directed RNA polymerase subunit beta, rpoB | 9 | 11 | 0.047 |
| CTL0568 (CT316) | 50S ribosomal protein L7/L12, rplL | 2 | 20 | 0.209 |
| CTL0570 (CT318) | 50S ribosomal protein L1, rplA | 4 | 18 | 0.049 |
| CTL0595 (CT341) | Chaperone protein, dnaJ | 3 | 11 | 0.003 |
| CTL0655 (CT398) | Hypothetical Protein | 3 | 19 | 0.013 |
| CTL0679 (CT421.1) | Hypothetical Protein | 3 | 30 | 0.699 |
| CTL0696 (CT437) | Elongation factor G, fusA | 19 | 32 | 0.132 |
| CTL0752 (CT491) | Transcription termination factor, rho | 4 | 11 | 0.040 |
| CTL0769 (CT507) | DNA-directed RNA polymerase subunit alpha, rpoA | 4 | 18 | 0.052 |
| CTL0770 (CT508) | 30S ribosomal protein S11, rpsK | 3 | 31 | 0.907 |
| CTL0771 (CT509) | 30S ribosomal protein S13, rpsM | 4 | 35 | 0.325 |
| CTL0774 (CT512) | 30S ribosomal protein S5, rpsE | 2 | 10 | 0.156 |
| CTL0776 (CT514) | 50S ribosomal protein L6, rplF | 3 | 25 | 0.629 |
| CTL0786 (CT524) | 30S ribosomal protein S19, rpsS | 2 | 14 | 0.759 |
| CTL0788 (CT526) | 50S ribosomal protein L23, rplW | 3 | 45 | 0.300 |
| CTL0789 (CT527) | 50S ribosomal protein L4, rplD | 4 | 24 | 0.018 |
| CTL0791 (CT529) | Putative inclusion membrane protein | 3 | 20 | 0.970 |
| CTL0863 (CT600) | Peptidoglycan-associated lipoprotein, pal | 2 | 15 | 0.509 |
| CTL0866 (CT603) | Thioredoxin peroxidase, ahpC | 11 | 74 | 0.280 |
| CTL0872 (CT608) | ATP-dependent DNA helicase, uvrD | 2 | 6 | 0.203 |
| CTL0874 (CT610) | Virulence protein, CADD | 5 | 44 | 0.259 |
| CTL0882 (CT618) | Putative inclusion membrane protein | 4 | 29 | 0.817 |
| CTL0018 (CT650) | Recombinase RecA | 3 | 10 | 0.021 |
| CTL0024 (CT655) | 2-dehydro-3-deoxyphosphooctonate aldolase, kdsA | 3 | 25 | 0.012 |
| CTL0046 (CT677) | Ribosome recycling factor, frr | 2 | 20 | 0.043 |
| CTL0060 (CT691) | Hypothetical Protein | 2 | 8 | 0.072 |
| CTL0112 (CT743) | Histone H1-like protein, hctA | 2 | 14 | 0.332 |
| CTL0137 (CT768) | Hypothetical Protein | 4 | 8 | 0.006 |
| CTL0140 (CT771) | Hydrolase | 3 | 25 | 0.080 |
| CTL0142 (CT773) | Leucine dehydrogenase, ldh | 4 | 18 | 0.009 |
| CTL0168 (CT799) | 50S ribosomal protein L25, rplY | 5 | 21 | 0.287 |
| CTL0183 (CT812) | Polymorphic membrane protein, pmpD | 6 | 7 | 0.009 |
| CTL0213 (CT841) | ATP-dependent zinc metalloprotease, ftsH | 3 | 9 | 0.0081 |
| CTL0233 (CT858) | Protease-like activity factor, CPAF | 2 | 5 | 0.914 |
| CTL0250 (CT871) | Polymorphic membrane protein, pmpG | 4 | 7 | 0.412 |
| CTL0254 (CT874) | Polymorphic membrane protein, pmpI | 3 | 4 | 0.011 |
| pL2-03 (pGP1) | Putative helicase | 2 | 7 | 0.038 |

### Infection with *C. trachomatis* alters the mitochondrial proteome to promote cell survival and mitochondrial fusion

Performing mass spectrometry on total protein content allowed for the comparison of host protein changes between mitochondria isolated from uninfected and infected cells. There was an average of 2,306 proteins identified per replicate with no significant difference between experimental replicates or between treatment conditions. When comparing crosslinked to non-crosslinked conditions, there were few differences in protein content with only 75 (2.8%) total proteins identified only in non-crosslinked conditions and 57 (2.1%) proteins only identified in the crosslinked conditions out of a total of 2,685 proteins in all replicates. Of all proteins identified, over 20% were known mitochondrial proteins (Figure 4A). Forty percent of proteins identified were cytoplasmic proteins and likely interact with mitochondrial proteins. One concern with isolating mitochondria is the close association with other organelles, particularly the endoplasmic reticulum and Golgi-derived vesicles. Within the proteomes, around 8% and 5% of proteins were known to interact with the ER or Golgi/Golgi-derived vesicles respectively, indicated that there was minimal pulldown of these organelles with the isolated mitochondria. Less than 2% of the proteins identified are predicted to be localized to the lysosomes, peroxisomes, or cytoskeletal elements. Proteins known to localize with these non-mitochondrial organelles may indicate direct interactions between mitochondria and non-mitochondrial organelles.

**Figure 4.**
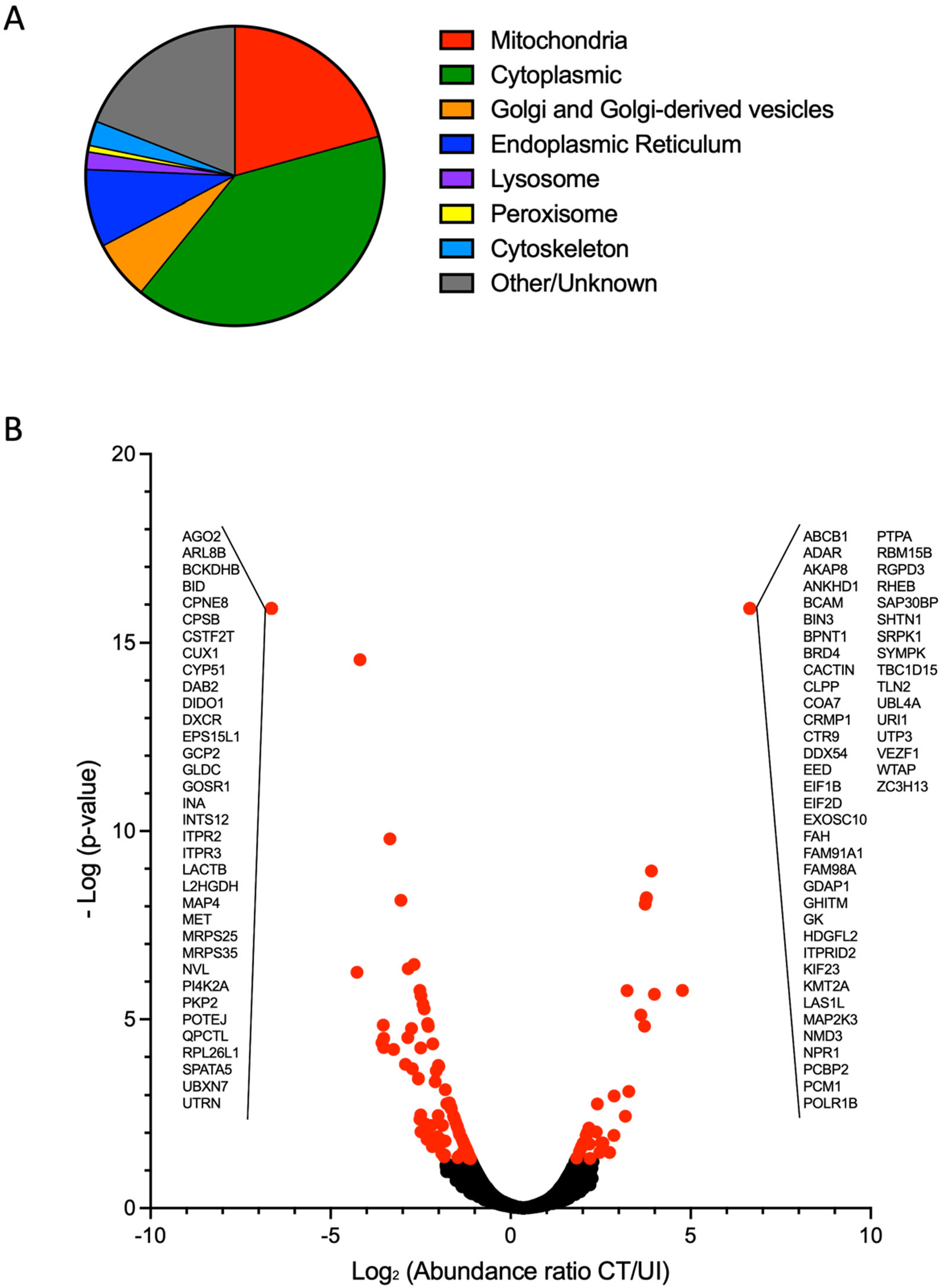
A) Gene Ontology Cellular Compartments for proteins identified in mass spectrometry screen of mitochondria from uninfected and infected cells (GO:0005739, 0005759, 0005743, 0005741, 0005758, 0097679, 0000139, 0044233, 0005783, 0032865, 0015629, 0005764, 0005777). B) Host protein peak abundances for each identified peptide were compared between groups and reported as log2 of the ratio of CT to UI where significant changes (red) were determined at p<0.05. All peptides identified in only one condition have the same value on the plot and are listed in insets.

To determine mitochondrial proteome differences between *C. trachomatis* infected and uninfected conditions, an abundance ratio was calculated by dividing the abundance of protein in the infected condition by the uninfected condition (CT/UI). There were 83 hits with significantly (p>0.05) increased protein abundance in infected conditions (Supplementary Table 1) and 125 hits that were significantly decreased in infected conditions (Supplementary Table 2) totaling 208 (7.8%) proteins with differential abundance. (Figure 4A). Of the proteins with differential abundance, there were a subset of proteins identified which were unique to one condition. Thirty-five proteins were present only in mitochondria of uninfected cells and 50 proteins were present only in mitochondria of infected cells (Figure 5).

**Figure 5.**
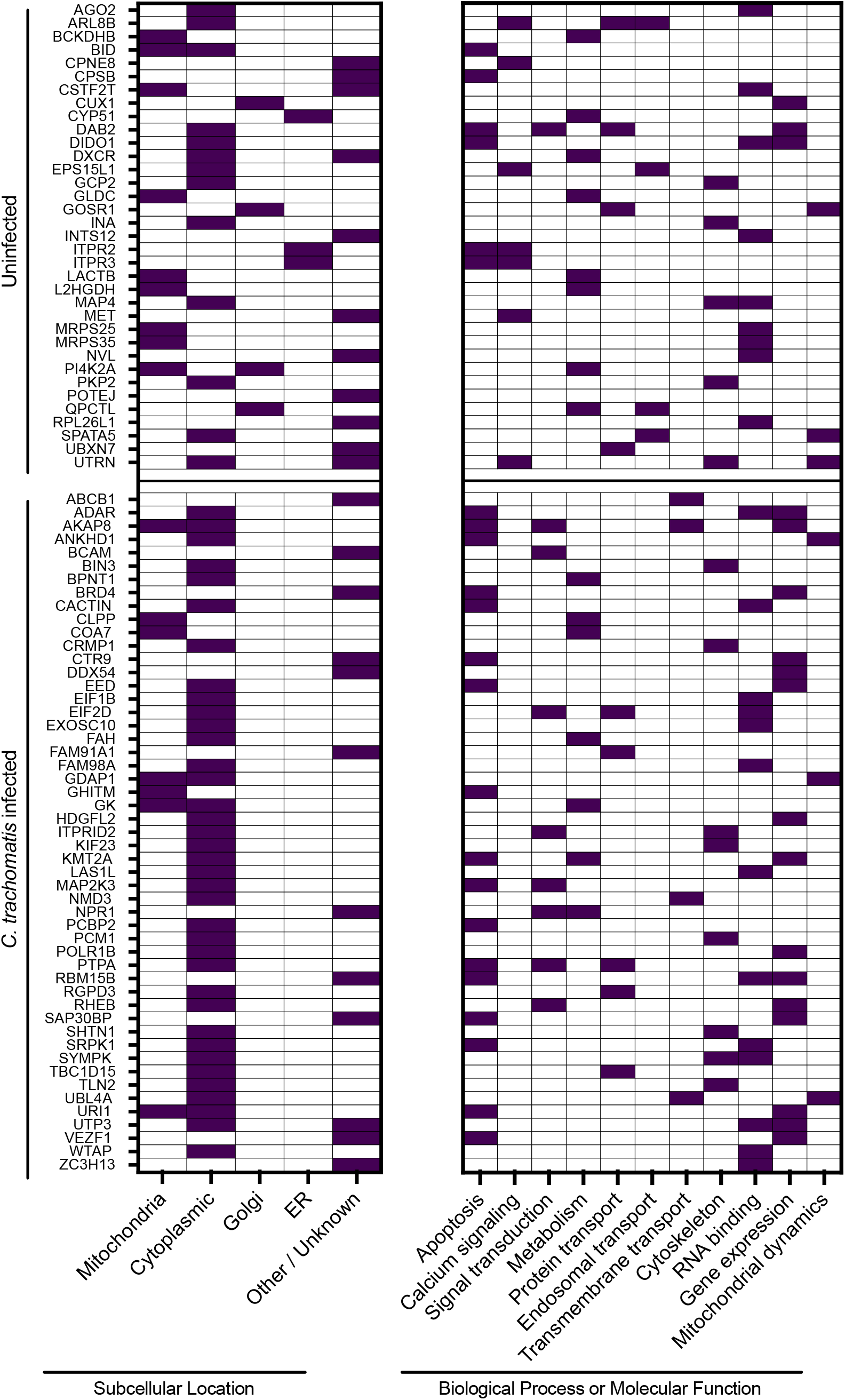
Mitochondria from uninfected (UI) or *C. trachomatis* L2-infected (CT) HeLa cells have unique protein content. Proteins unique to one condition (uninfected or *C. trachomatis* infected). Known or estimated subcellular locations based on Gene Ontology Cellular Compartments (GO: 0005739, 0005759, 0005743, 0005741, 0005758, 0097679, 0000139, 0044233, 0005783, 0032865) and functions based on Gene Ontology Biological Process or Molecular Function (GO:0006915, 0043065, 0043066, 0045087, 0019722, 0050848, 0007165, 0005975, 0008610, 0006633, 0046112, 0009058, 0015031, 0016197, 0030036, 0005200, 0003723, 1903108, 0003677, 0006306, 0000266, 0008053) are indicated by filled in squares.

Proteins present at the mitochondria only of uninfected cells may be recruited elsewhere during infection. Specifically, infection restricted the presence of several Golgi and ER proteins. Functional analysis of proteins unique to the uninfected condition showed that they were representative of diverse functions including RNA binding (26%), metabolism (23%), calcium signaling (20%), and apoptosis (17%). The proteins involved in apoptosis that were present only in the uninfected cell mitochondria were all pro-apoptotic effectors, including well-characterized BH3 domain Bid, which localizes to the mitochondrial membrane during initiation of intrinsic apoptosis [44]. Additionally, three positive regulators of mitochondrial fission were present only in uninfected cells.

Of those proteins present only in infected replicates, 37 are known cytoplasmic proteins, suggesting that chlamydial infection results in the relocation of these proteins to the mitochondria, potentially interacting with proteins on the mitochondrial membrane. There was a functional symmetry to those proteins unique to the infected condition and those absent during infection. Of the unique mitochondrial proteins in the infected cells, the majority were involved in apoptosis (34%), RNA binding (26%) and metabolism (14%) (Figure 5). However, in this condition, proteins involved in apoptosis act largely as anti-apoptotic factors and those involved in metabolism were oxidative phosphorylation and glycolysis activators.

## Discussion

In this study, we describe the identification of mitochondrial targeting sequences in the *C. trachomatis* genome and demonstrate that at least five of these signals are recognized and result in the translocation of these proteins to the mitochondria in HeLa cells. Previous work has shown that knock down of components of the Tom complex are essential for replication of *C. caviae* either in *Drosophila* cells or mammalian cells [45]. The Tom complex is a multiprotein complex involved in the recognition and import of nuclear encoded mitochondrial proteins to mitochondria. This requirement for components of the Tom complex, Tom40 and Tom22, was specific for *C. caviae*, however, as knockdown of Tom40 and Tom22 were not inhibitory to *C. trachomatis* [45]. It is unclear which, if any, *C. caviae* proteins might be trafficked to mitochondria or whether *C. trachomatis* mitochondrially targeted proteins might utilize different mechanisms. The chlamydial effectors which co-localized with mitochondria are the first reports of *C. trachomatis* proteins which may directly interact with host mitochondria. From the initial bioinformatic screen, eight proteins were not tested in the ectopic expression analysis. While those with *C. trachomatis* L2 orthologs were not later identified in the mitochondria of infected cells, it remains possible that the three proteins unique to serovar D, CT166, CT168 and CT352, may also be mitochondria-targeting effectors.

Examination of the MTS for the five positive colocalizing proteins after ectopic expression reveals that there may be a preference for an MTS around 35 amino acids in length. Only CT642 had an MTS that was outside of the 32-37 amino acid range and interestingly, this MTS was one of the shortest detected by MitoProt at 8 amino acids. Despite being almost a quarter the length of the other sequences, the CT642 MTS had an equal ratio of conserved residues (Figure 1B, underlined) which likely contribute to the ability to be recognized by cognate machinery. Cleavage sequences were identified for only two of the five proteins, CT618 and CT647. It remains unclear whether these signals are cleaved or if there are alternate cleavage mechanisms for the other MTS. Adding evidence to the functionality of the MTS signal, deleting the sequence from the GFP-fusion proteins leads to loss of mitochondrial localization. Both CT529 and CT642 localized to discrete subcellular compartments rather than a diffuse cytoplasmic staining. This phenotype may be due to their Inc-like structure as it has been shown previously that ectopically expressed Incs cause the formation of or associate with distinct vesicles within HeLa cells [46].

Because CT529, CT618 and CT642 have been previously identified as putative Incs [42], it was not surprising that each were confirmed as Type 3 secreted by heterologous *Y. pseudotuberculosis* system (Figure 3). Within *Chlamydia* species, both CT618 and CT642 appear to be conserved [47], so it is possible that the MTS is conserved as well. CT132 and CT647 were not predicted to be T3S, however, it is possible that these proteins could be secreted through the general secretory system.

The proteomics analysis of isolated mitochondria was critical in demonstrating the validity of the mitochondrial localization for CT529 and CT618 from the initial screen as both were included in the 49 chlamydial proteins identified (Table 2). Both CT529 and CT618 were previously identified in a T3S component screen as interacting with the same unique T3S chaperone, CT260 (Mcsc, <u>M</u>ultiple <u>c</u>argo <u>s</u>ecretion <u>c</u>haperone) [35]. This chaperone may be important for proper secretion of CT529 and CT618 as they are translocated through the T3SS and recognized by host chaperone proteins. CT529 has been previously demonstrated to colocalize with lipid droplets, however, this study artificially stimulated lipid droplet formation, which may not reflect the localization of this protein during a normal infection of HeLa cells [10]. Additionally, when CT529 was expressed with a C-terminal GFP, there did appear to be protein predominately localized outside of lipid droplets, likely mitochondria considering the data presented herein. This localization was absent when CT529 was expressed with an N-terminal mCherry, which likely disrupted the MTS, inhibiting the normal translocation of this effector. The function of CT529 and CT618 at the mitochondria remains unknown and additional studies will be needed to determine the role these proteins play during chlamydial infection.

In the initial Mitoprot MTS screen, we were only able to identify N-terminal MTS sequences. While many mitochondrial proteins rely on an N-terminal MTS signal, there are both internal MTS [48] and unique non-N-terminal MTS in β-barrel proteins [49]. Therefore, the proteomics screen enabled us to identify additional chlamydial candidate effectors which may localize to mitochondria during infection (Table 2). It is possible that these chlamydial effectors may traffic to the mitochondria through alternate MTS, or through direct or indirect proteins which associate with the mitochondrial outer membrane. While the majority of chlamydial proteins identified at the mitochondria of infected cells may have been surprising, two proteins which were likely to interact with mitochondrial proteins were identified; CT858 (CPAF) and CT610 (CADD). CPAF (<u>C</u>hlamydial <u>p</u>rotease-like <u>a</u>ctivity <u>f</u>actor) has been hypothesized and demonstrated to have a variety of functions, particularly in cleaving a set of host and chlamydial proteins [50]. Pirbhai et al provided evidence that CPAF may cleave BH3 family proteins (like Bid) during chlamydial infection to prevent apoptosis [51] and in a study in Protochlamydia, a CPAF homolog was shown to disrupt mitochondrial membrane potential [52]. These studies suggest that CPAF may target proteins of the mitochondria, which is supported by its identification in the mitochondrial proteome here. In fact, absence of Bid at the mitochondria of infected cells (Figure 5B) could be explained by its cleavage by CPAF. While the protein targets of CPAF have been disputed, Bid cleavage has not been specifically examined *in vivo* [53]. CADD (Chlamydia protein associating with death domains) has shown to be involved in the apoptotic pathway and although the initial analysis of CADD did not detect CADD localization at mitochondria with CADD antiserum [54], our data suggests that CADD may interact with mitochondrially localized apoptotic effectors. Our initial bioinformatic screen did not identify CT421.1 as a candidate MTS, as the MitoProt probability fell just below the cutoff of 0.7. However, this protein was identified in all replicates of mitochondria from infected cells and it is possible that CT421.1 does have a functional MTS. Additional studies will be needed to investigate the validity of this promising candidate.

Through the proteomic screen, we were able to demonstrate changes in the protein content of mitochondria during infection (Figure 4A). Infection with *C. trachomatis* resulted in an absence of several pro-apoptotic factors and recruitment of anti-apoptotic factors. One regulatory pathway of apoptosis that appeared to be disproportionately affected was the mTORC1 (<u>m</u>olecular <u>t</u>arget of <u>r</u>apamycin <u>c</u>omplex 1) pathway. The master activator of this pathway, RheB, was present only in infected mitochondria, suggesting activation of this pathway by chlamydia [55]. Along with RheB, the downstream mTORC1 effector, URI1, which maintains phosphorylation and subsequent inactivation of Bad, a pro-apoptotic Bcl-2 family protein was present at mitochondria only in infected cells[56]. Several putative negative regulators of mTORC1 were also absent from or decreased in abundance in mitochondria of infected cells, including NRDC (nardilysin) [57]. Another apoptotic pathway affected by infection included the calcium influx into the mitochondria by the endoplasmic reticulum. Inositol 1,4,5-triphosphate receptor interacts with the mitochondrial outer membrane porin, VDAC (<u>v</u>oltage-<u>d</u>ependent <u>a</u>nion <u>c</u>hannel type 1), resulting in Ca^2+^ efflux into the mitochondria [58]. The inositol 1,4,5-triphosphate receptor(s) are known to be recruited to the inclusion membrane by microdomain Inc, MrcA, which may explain their absence in the infected proteome [59, 60]. This report contributes to a growing body of knowledge within the field detailing the variety of mechanisms *Chlamydia* spp. utilize to inhibit apoptosis [61–64].

In addition to changes to apoptotic pathways, *C. trachomatis* infection resulted in a shift of mitochondrial dynamics toward promotion of mitochondrial fusion, which corroborates similar findings [22, 63, 65, 66]. These changes were identified by the absence of pro-fission regulator UTRN (utrophin) [67] and presence of pro-fusion regulators ANKHD1 (Ankyrin repeat and KH domain-containing protein 1) [68], and UBL4A (Ubiquitin-like protein 4A) [69] only in infected cells (Figure 5). Mitochondrial dynamics play a critical role in metabolism and fusion has been demonstrated to support increased oxidative phosphorylation, and production of ATP [70]. Additionally, there were changes in the mitochondrial proteome to reflect this shift in increased metabolism. Particularly, glycerol kinase (GK) which has been suggested to bind to the mitochondrial porin, during increased metabolic activity [71], was bound to mitochondria only in infected cells. The proteomics data presented here suggests shifts in mitochondrial function during infection and can serve as a resource to further research into the specific changes induced by *C. trachomatis*.

In summary, we demonstrate that *C. trachomatis* secretes at least two proteins which associate with host mitochondria. The individual and collective roles of these proteins remain to be resolved, but it is clear that chlamydial infection induces profound changes in mitochondrial proteome composition. An improved understanding of how *Chlamydia* manipulate mitochondria and other cellular organelles should provide new insights into how intracellular pathogens create favorable intracellular niches for replication.

## Acknowledgements

This work was supported by the Intramural Research Program of the NIAID, NIH. We thank Rebecca Miller for technical assistance and Adam Nock for review of this manuscript.

---

**Supplemental Figure 1.**
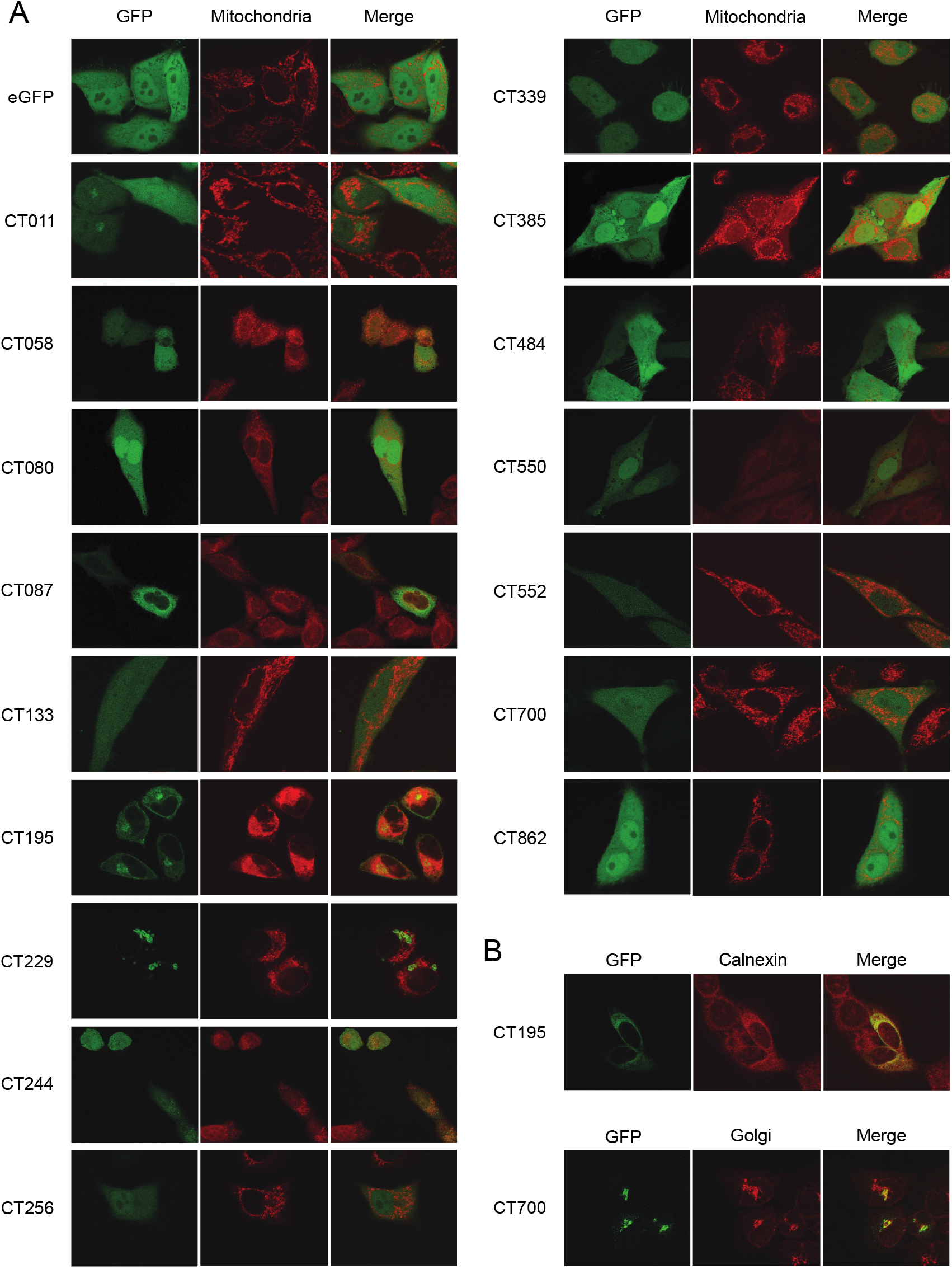
A) Immunofluorescent staining shows GFP localization of transfected *C. trachomatis* proteins which did not have functional MTS. HeLa cells on coverslips were transfected with either the empty EGFP-N1 vector (eGFP) or GFP-tagged *C. trachomatis* proteins (CT###) and incubated for 24 hours before staining with 100nM MitoTracker (red). B) Colocalizations with other organelles demonstrated with anti-calnexin antibodies or anti-Golgi antibodies. Bar = 10 μm.

**Supplemental Table 1.**
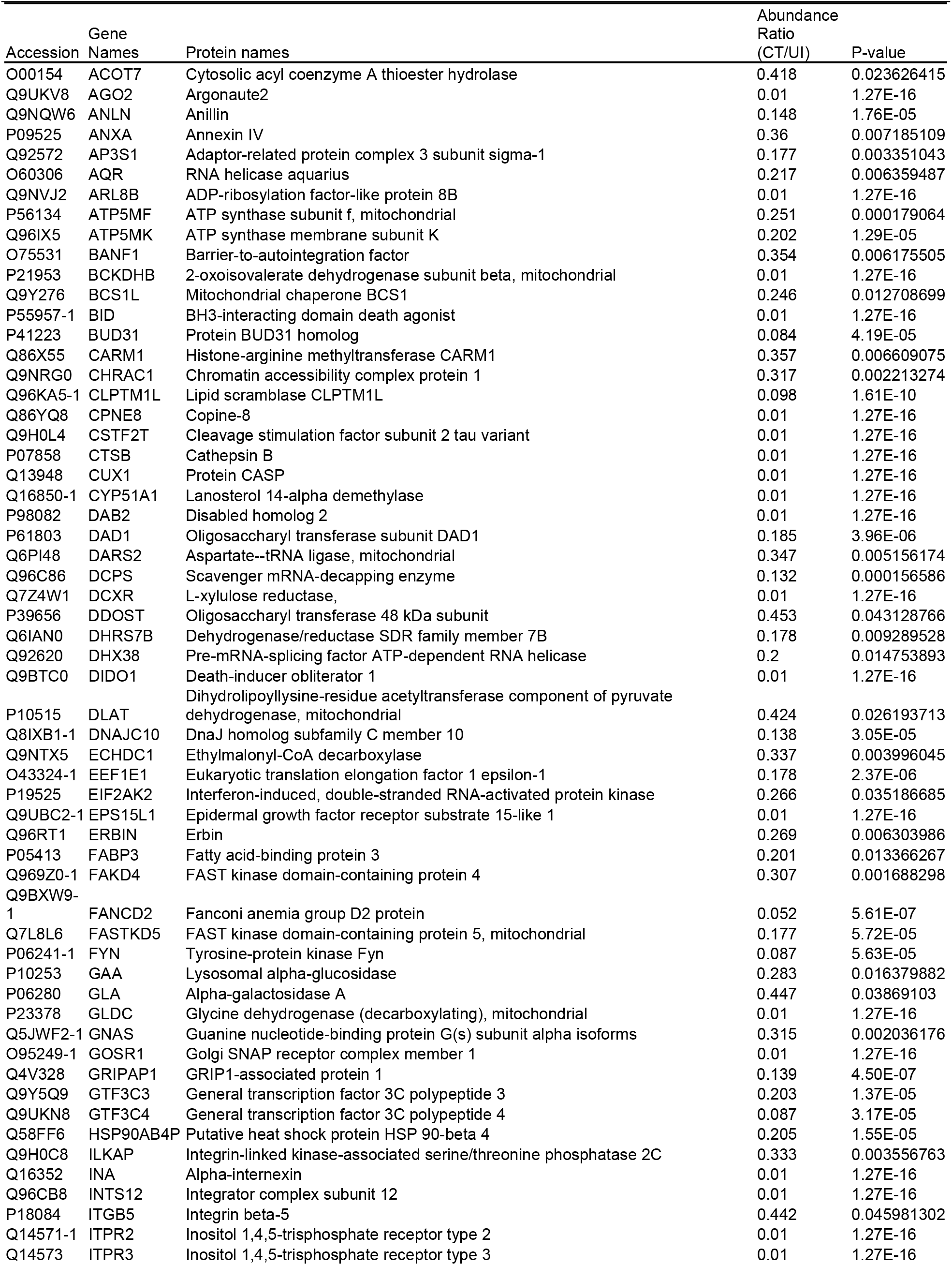
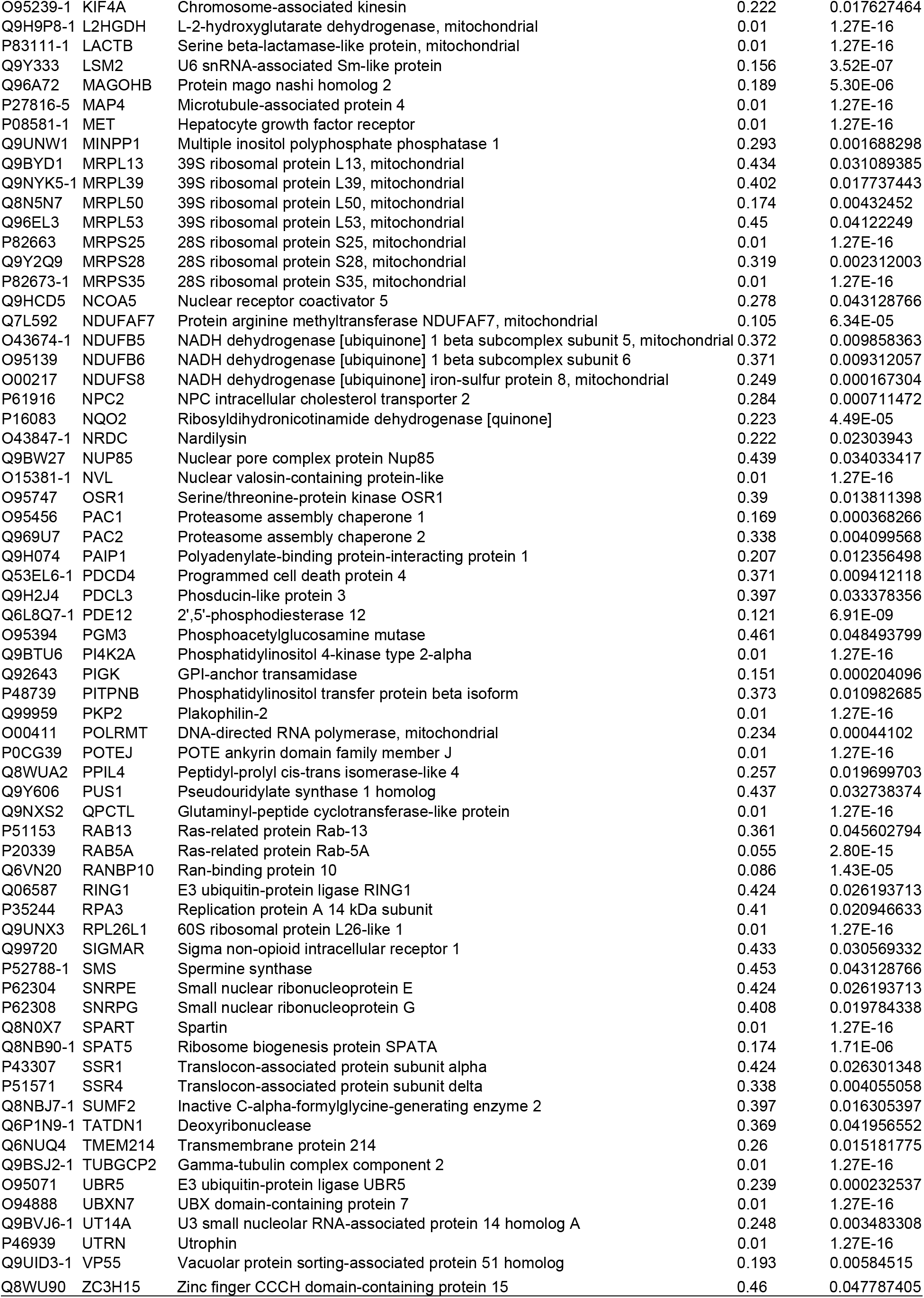
Proteins with decreased abundance in mitochondria of infected cells

**Supplemental Table 2.**
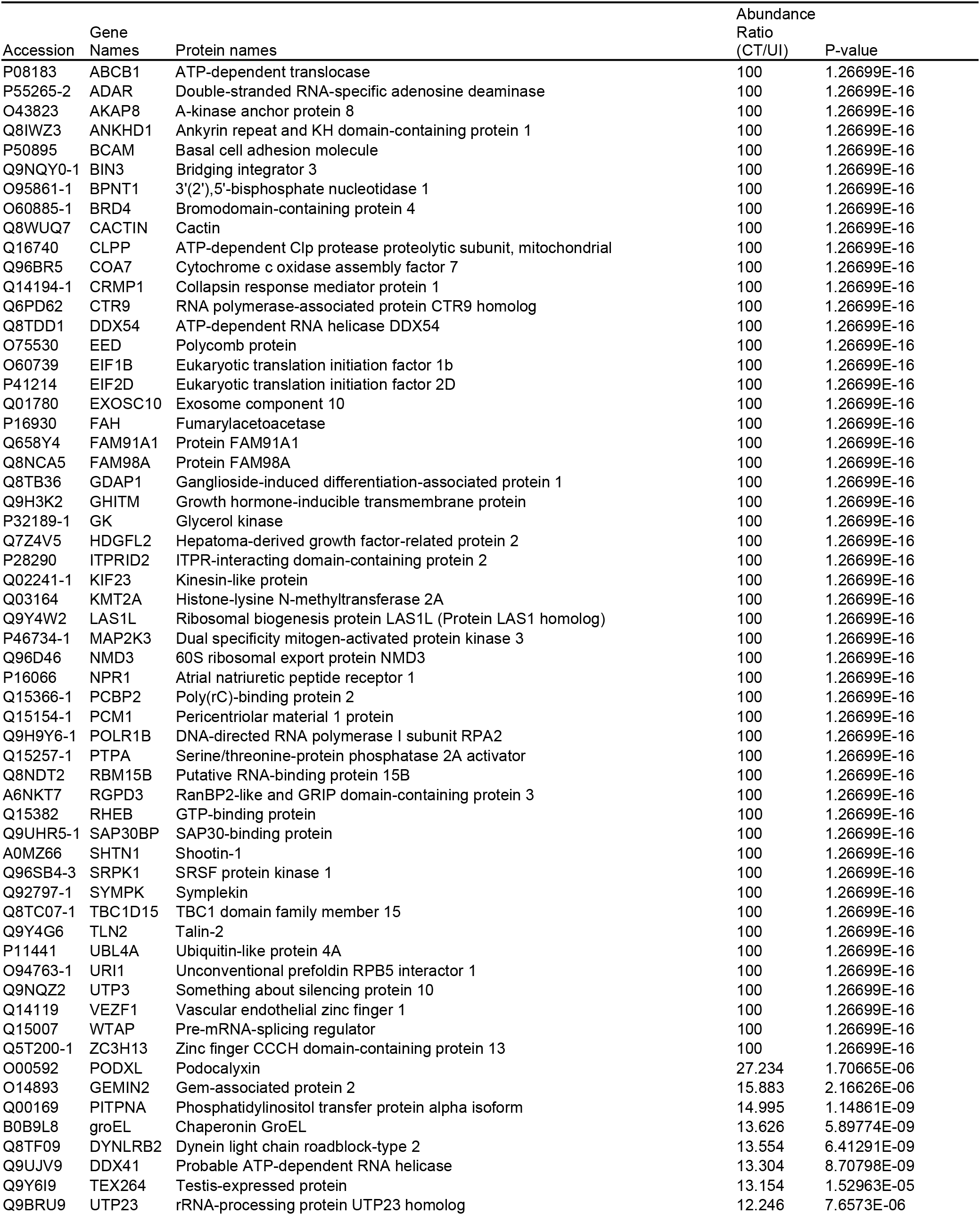
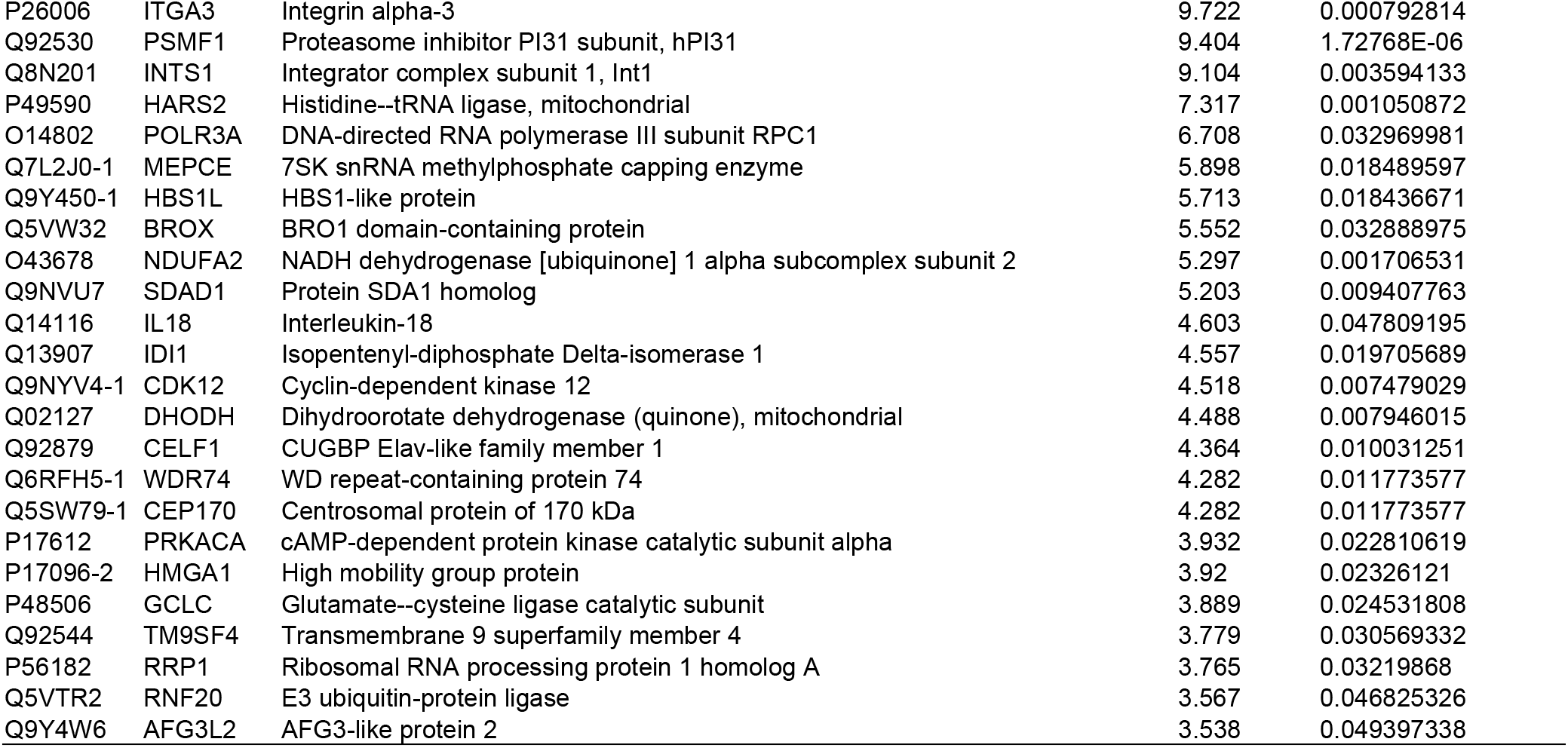
Proteins with increased abundance in mitochondria of infected cells

Supplemental Table 3. Proteome Dataset from LC/MS (Excel file)

